# Building model prototypes from time-course data

**DOI:** 10.1101/2022.01.27.478080

**Authors:** Alan Veliz-Cuba, Stephen Randal Voss, David Murrugarra

## Abstract

A primary challenge in building predictive models from temporal data is selecting the appropriate model topology and the regulatory functions that describe the data. Software packages are available for equation learning of continuous models, but not for discrete models. In this paper we introduce a method for building model prototypes. These model prototypes consist of a wiring diagram and a set of discrete functions that can explain the time course data. The method takes as input a collection of time course data or discretized measurements over time. After network inference, we use our toolbox to simulate the prototype model as a stochastic Boolean model. Our method provides a model that can qualitatively reproduce the patterns of the original data and can further be used for model analysis, making predictions, and designing interventions. We applied our method to a time-course, gene-expression data that were collected during salamander tail regeneration under control and intervention conditions. The inferred model captures important regulations that were previously validated in the research literature and gives novel interactions for future testing. The toolbox for inference and simulations is freely available at github.com/alanavc/prototype-model.

## 1 Introduction

The process of constructing discrete models from experimental data has several steps that have been studied in parallel. The main steps involved in this process are discretization, network inference, network selection, model interpolation, and deterministic/stochastic simulations [1–8]. Although some tools exist that address the global process [9–11], either the code is unavailable, not editable, or not in a ready-to-use format.

Equation learning (EQ) methods for differential equation (DE) models start with a collection of time course data and then “recovers” the governing equations using a library of functions [12,13]. Many methods for EQ of DE models are based on formulating the inference problem as a parameter estimation problem that can be solved via optimization techniques [12,13]. Analogue methods for equation learning of discrete models that can learn both the network topology and the functions are still under development. Some of these existing methods can provide network candidates (i.e., possible wiring diagrams) that can explain the data. Other methods can provide candidate functions based on interpolating the data.

The main contribution of the paper is the combination of methods and the concrete toolbox that any user can use without familiarity with algebraic techniques. Our toolbox is modular, so that any step in the flowchart can be modified by the user without any restrictions. Importantly, it is also open-source and is freely available through a GitHub site. It works in Octave, so it is available for use in any operating system without the need of any license costs due to proprietary software. This makes our results fully reproducible.

The starting point of our method is experimental time-course data. Our focus is the construction of Boolean models, but we show with a toy model how our method also works for mixed-state models where variables can have different number of states or levels. As an application, we construct a model prototype using gene expression data for several time points which was collected during tail regeneration experiments in axolotls. We also use a synthetic network to illustrate the effect of data size, noise, and number of levels.

## 2 Methods

We will use “network” to refer to the correct wiring diagram and set of functions, and “model” to refer to the wiring diagram and set of functions that are obtained using data generated by the network.

Here we describe the methods for model selection (i.e., wiring diagram and regulatory functions) and the framework for simulations. We assume that we are given time courses of the form *s*^1^ → *s*^2^ → … → *s^r^*, where 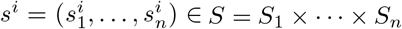. Here *S_i_* is a finite set of all the values that the i-th variable can take. Note that if *S_i_* = {0,1}, then we have a Boolean model.

### Example 2.1.

*To illustrate the methods, we use an example with the following four time courses.*

1. (0.1,1.1,1.9,0.9, 0.2) → (0.0,0.2,0.2,0.1,0.1) → (0.0,1.1,0.1,1.9, 2.1)
2. (1.9,0.1, 0.9,0.1, 0.0) → (0.9,1.1,0.1,1.9, 2.1) → (1.1, 0.9,0.1,1.9, 2.0)
3. (0.2,1.1,1.9, 0.9,1.1) → (0.1, 0.0,0.2, 0.1,0.1)
4. (0.1, 0.9, 2.1,1.1, 2.1) → (0.2, 0.1,0.2, 0.1,1.1)

### 2.1 Discretization

We implemented a simple discretization method based on binning data by dividing the range of the data into equally spaced regions. The time courses suggest that the number of levels for variables *x*_1_, *x*_2_, *x*_3_, *x*_4_, *x*_5_, are 3, 2, 3, 3, 3, respectively. For example, by plotting the values of *x*_1_ and *x*_2_ for each trajectory (Fig. 2), we see that *x*_1_ has 3 distinctive levels and *x*_2_ has 2 distinctive levels. For *x*_1_, all values below the dotted line will be mapped to 0 (low); all values between the dotted and dashed lines will get mapped to 1 (medium); and all values above the dashed line will get mapped to 2 (high). For *x*_2_, all values below the dotted line will be mapped to 0 (low); and all values above the dotted line will get mapped to 1 (high).

**Figure 1:**
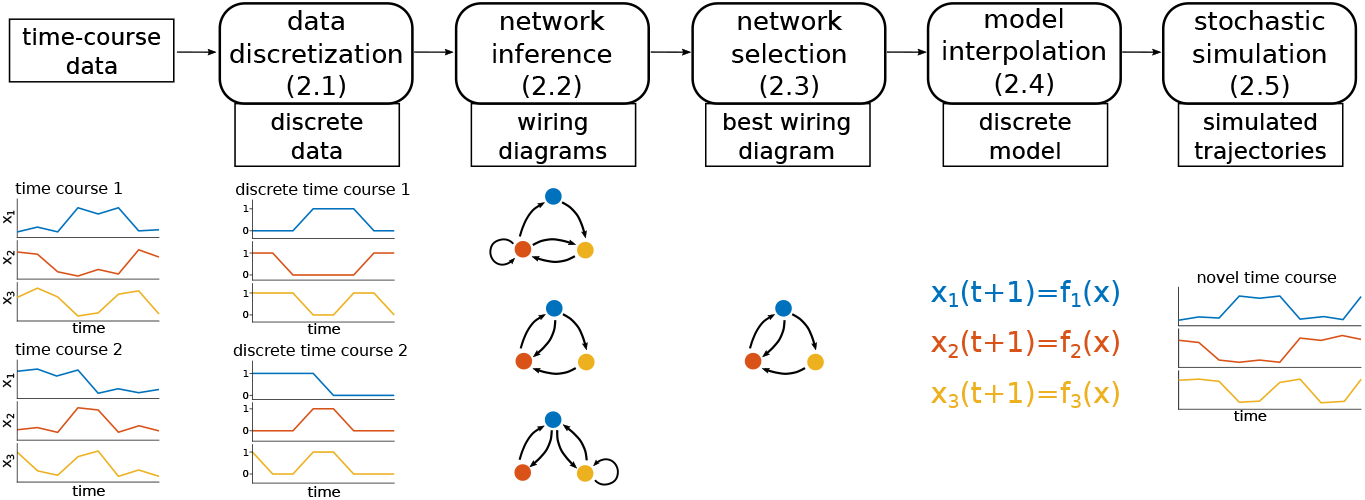
Flowchart showing the steps in model creation from data and the sections where each step is described. Starting from experimental time courses, we first transform the data into discrete values (in this case Boolean). Using algebraic techniques, we find wiring diagrams that explain the data. Each wiring diagram found will be consistent with all discrete time courses. We select the best wiring diagram and then find a discrete model that fits all the discrete data. This will result in a discrete model that can be simulated and compared with the original data. The model can also be run with new initial conditions or for longer time to create novel time courses that can be used to make predictions.

**Figure 2:**
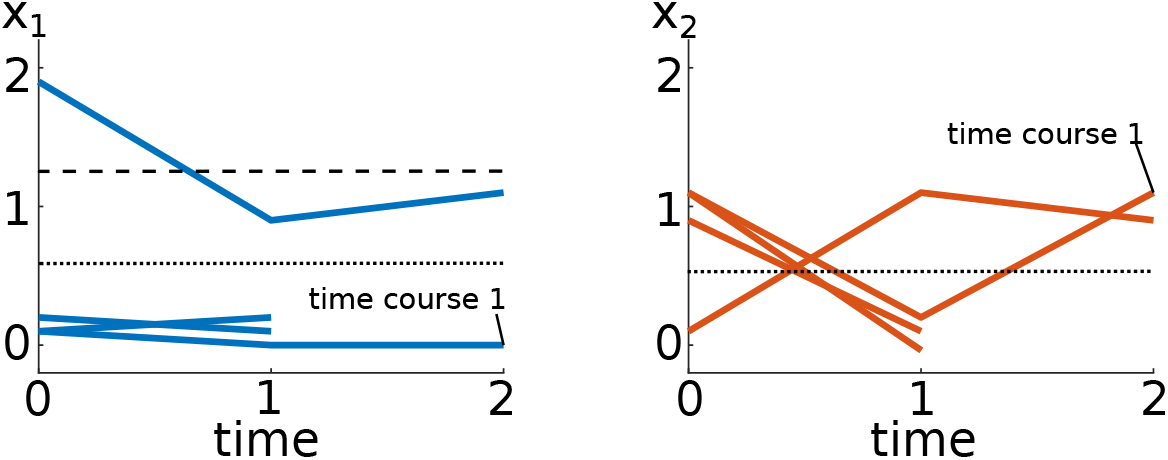
Values of *x*_1_ and *x*_2_ for the time courses. Variable *x*_1_ can be considered as having 3 levels, whereas variable *x*_2_ has 2 levels. The dashed lines show how the range of the data can be divided into regions (3 regions for *x*_1_ and 2 for *x*_2_), which will determine the discretization.

Then, the discrete time courses are given below.

1. 01210 → 00000 → 01022
2. 20100 → 11022 → 11022
3. 01211 → 00000
4. 01212 → 00001

In this case S = {0,1, 2} × {0,1} × {0,1, 2} × {0,1, 2} × {0,1, 2}.

### 2.2 Network Inference

To find the wiring diagrams that are consistent with a collection of time courses of the form *s*^1^ → *s*^2^ → 0.. → *s^r^* we use the algebraic framework introduced in [3]. This framework takes partial information about the evolution of a network *s* → *f* (*s*) and returns all the minimal wiring diagrams that are consistent with the data. This approach guarantees that for each minimal wiring diagram there exists a model that fits the data such that each interaction is activation or inhibition [3].

To use the framework in [3], we first note that each time course *s*^1^ → *s*^2^ → … → *s^r^* implies that *s*^*j*+1^ = *f* (*s^j^*) for *j* = 1,…,*r* — 1, where f is the network one is trying to infer. This results in a set *D* ⊆ *S* such that *f* (*s*) is known for every *s* ∈ *D*. That is, *D* is the set of inputs for which we know the outputs.

#### Example 2.2.

*In Example 2.1, D* = {01210,00000, 20100,11022, 01211, 01212}. *Then, the partial information we have is given by the table*

*Then, using the algebraic techniques in [3] we can find all minimal wiring diagrams that are consistent with the data. For each variable x_i_ in the network, the algebraic framework returns W*_1_, …, *W_k_, where each W_j_ is a minimal set of inputs for variable i. For our example we obtain Table 2.*

**Table 1:** Partial information for example.

| $x$ | $f(x)$ |
| --- | --- |
| 01210 | 00000 |
| 00000 | 01022 |
| 20100 | 11022 |
| 11022 | 11022 |
| 01211 | 00000 |
| 01212 | 00001 |

**Table 2:** Minimal wiring diagrams. The +/- superscripts indicate activation/inhibition. For example, the set 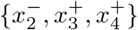 indicates that one way to explain the data is for *x*_2_ to be an inhibitor of *x*_1_ and *x*_3_ and *x*_4_ to be activators of *x*_1_. By choosing one set for each variable, one obtains a wiring diagram that is consistent with the data. Note that no variable affects *x*_3_ (i.e., constant function).

| $x_i$ | minimal wiring diagrams for $x_i$ |
| --- | --- |
| $x_1$ | $\{x_1^+\}, \{x_2^-, x_3^+, x_4^+\}$ |
| $x_2$ | $\{x_3^-\}, \{x_2^-, x_4^+\}, \{x_1^+, x_4^-\}, \{x_1^+, x_2^-\}$ |
| $x_3$ | $\{\}$ |
| $x_4$ | $\{x_3^-\}, \{x_2^-, x_4^+\}, \{x_1^+, x_4^-\}, \{x_1^+, x_2^-\}$ |
| $x_5$ | $\{x_3^-, x_5^+\}, \{x_2^-, x_4^+, x_5^+\}, \{x_1^+, x_4^-, x_5^+\}, \{x_1^+, x_2^-, x_5^+\}$ |

By selecting one wiring diagram for each *x_i_*, we obtain a (global) wiring diagram that is consistent with the data. For example, if we select 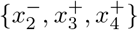 for 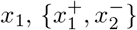 for *x*_2_, { } for *x*_3_, 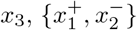 for *x*_4_, and 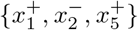 for *x*_5_, we obtain the wiring diagram shown in Fig. 3. To compare different wiring diagrams we can use the adjacency matrix representation.

**Figure 3:**
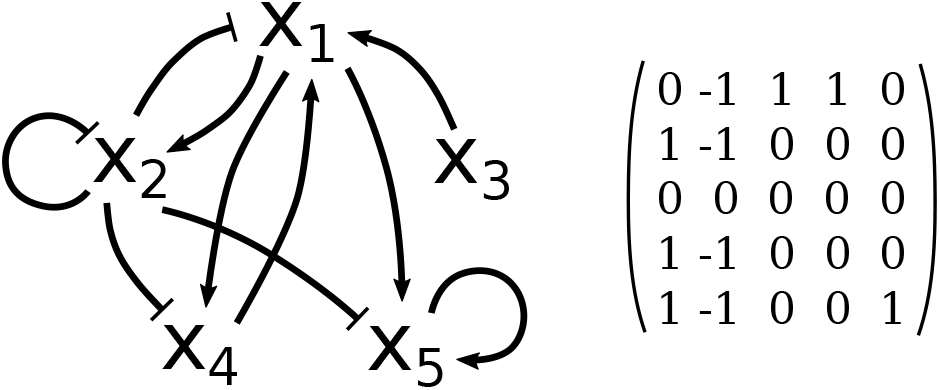
Example of wiring diagram consistent with the data. Left: Wiring diagram. Right: Adjacency matrix representation.

### 2.3 Wiring diagram selection

The network inference described in Section 2.2 could return several minimal network candidates for each variable. That is, for a given time course data, there might be several models that explain the data and that are minimal. The method will return all candidate wiring diagrams. In order to select one model out of all possible options, we calculate the “best wiring diagram” by including only the most frequent interactions from the wiring diagrams found. For each variable, *x_i_*, we quantified the frequency 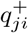 of positive interactions *x_j_*→ *x_i_* across all possible wiring diagrams and the frequency 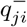 of negative interaction 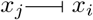 across all possible wiring diagrams for all *j* = 1,…, *n*. That is, the parameter 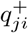 (resp. 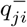) represents the frequency of regulator 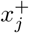 (resp. 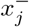) in the minimal wiring diagrams of *x_i_* (see Example 2.3 and Table 3 for additional details). Then we construct an adjacency matrix *W*\* by considering the interactions with a frequency above certain threshold *τ*. If conflicts arise (that is, when 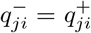 for some *j*), then we discard those interactions. Subsequently, for each row of *W*\*, say 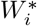, we calculate the distance with each possible wiring diagram of *x_i_* (these are represented as rows). Finally, we construct an adjacency matrix W with rows corresponding to the rows with minimum distances.

**Table 3:** Frequencies of interactions on minimal wiring diagrams. The parameter 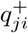 (resp. 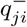) represents the frequency of regulator 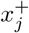 (resp. 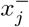) in the minimal wiring diagrams of *x_i_*. For instance, for variable *x*_1_ in the first row, 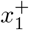 appears in one out of two wiring diagrams, therefore 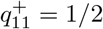.

|  | Frequencies of activations and inhibitions | # of WDs |
| --- | --- | --- |
| $x_1$ | $q_{11}^+ = 1/2, q_{21}^- = 1/2, q_{31}^+ = 1/2, q_{41}^+ = 1/2$ | 2 |
| $x_2$ | $q_{12}^+ = 2/4, q_{22}^- = 2/4, q_{32}^- = 1/4, q_{42}^+ = 1/4, q_{42}^- = 1/4$ | 4 |
| $x_3$ | NA | 0 |
| $x_4$ | $q_{14}^+ = 2/4, q_{24}^- = 2/4, q_{34}^- = 1/4, q_{44}^- = 1/4, q_{44}^+ = 1/4$ | 4 |
| $x_5$ | $q_{15}^+ = 2/4, q_{25}^- = 2/4, q_{35}^- = 1/4, q_{45}^+ = 1/4, q_{45}^- = 1/4, q_{55}^+ = 4/4$ | 4 |

#### Example 2.3.

*For the network in Example 2.1, we calculated the frequency of the interactions; see Table 3. For instance, for variable x*_1_ *in the first row of Table 3,* 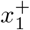 *appears in one out of two wiring diagrams (see Table 2), therefore* 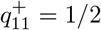. *Then, we computed an adjacency matrix W* by including the interactions with a frequency above the threshold τ* = 1/5. *We discarded conflicting interactions (i.e., the cases where* 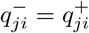).

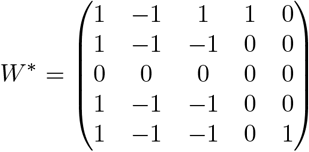

*Then, for each row of W*\*, *say* 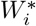, *we calculate the distance with each possible wiring diagram of x_i_ (these are represented as rows). Then we construct an adjacency matrix with rows corresponding to the rows with minimum distances. Then, the matrix after the distance calculations is*:

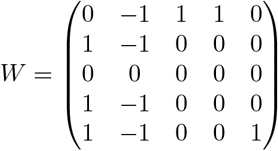

The reason for why we take a distance approach is because there might not be a truth table satisfying the matrix *W*\* but there is certainly one for *W* as shown in Example 2.2.

### 2.4 Fitting Model to Data

After one wiring diagram has been selected from the family of minimal wiring diagrams, we proceed to construct a function that fits the data. Although there are known formulas for interpolation, we are interested in *monotone* interpolation, that is, we need to find a model that not only fits the data, but one whose signs of interaction match the wiring diagram selected.

We illustrate our approach with wiring diagram 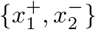 for variable *x*_4_. Since it is guaranteed that there is a monotone function *h*(*x*_1_,*x*_2_) that fits the data for variable *x*_4_ [3], we consider Table 1 with only *x*_1_ and *x*_2_ in the first column (inputs) and only *x*_4_ in the second column (output).

We now rewrite this table as a truth table by ordering the inputs lexicographically (*x*_1_ ∈ {0,1, 2}, *x*_2_ ∈ {0,1}), where some entries are unknown, Table 5.

**Table 4:**
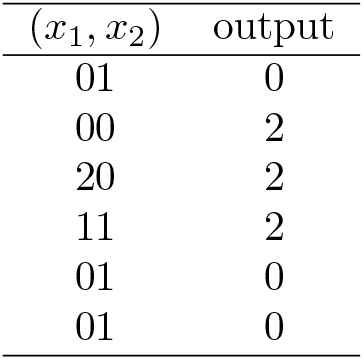
Partial information for variable *x*_4_ with wiring diagram 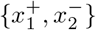.

**Table 5:** Incomplete truth table for variable *x*_4_ with wiring diagram 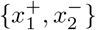 and the corresponding construction of a function for all inputs.

| $(x_1, x_2)$ | output | $h(x_1, x_2)$ |
| --- | --- | --- |
| 00 | 2 | 2 |
| 01 | 0 | 0 |
| 10 | ? | 2 |
| 11 | 2 | 2 |
| 20 | 2 | 2 |
| 21 | ? | 2 |

To fill in the table, we use the fact that the function increases with respect to *x*_1_ and decreases with respect to *x*_2_. For example, since *h*(2,1) ≥ *h*(1,1) = 2, it follows that *h*(2,1) = 2. Similarly, since 2 = *h*(0, 0) ≤ *h*(1, 0) ≤ *h*(2, 0) = 2, it follows that *h*(1, 0) = 2. In this way, we obtain the value of the missing entries. This process can be done for all wiring diagrams and for all variables. Since the existence of a wiring diagram guarantees that there is at least one suitable function that fits the data, this is always possible [3]. To guarantee that the fitting is unique, we implemented the algorithmic construction from Lemma 2.4 from [3].

### 2.5 Stochastic Framework

For the simulations we will use the stochastic framework introduced in [14] referred to as Stochastic Discrete Dynamical Systems (SDDS). This framework is a natural extension of Boolean networks and is an appropriate setup to model the effect of intrinsic noise on network dynamics. Consider the discrete variables *x*_1_,…, *x_n_* that can take values in finite sets *S*_1_,…, *S_n_*, respectively. Let *S* = *S*_1_ × … × *S_n_* be the Cartesian product. An *SDDS* in the variables *x*_1_,…, *x_n_* is a collection of n triplets

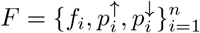

where

- *f_i_*: *S* → *S_i_* is the update function for *x_i_*, for all *i* = 1,…, *n*.
- 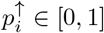 is the activation propensity.
- 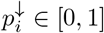 is the degradation propensity.

The stochasticity originates from the propensity parameters 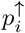 and 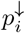, which should be interpreted as follows: If there would be an activation of *x_k_* at the next time step, i.e., if *s*_1_, *s*_2_ ∈ *S_k_* with *s*_1_ < *s*_2_ and *x_k_* (*t*) = *s*_1_, and *f_k_*(*x*_1_(*t*), …, *x_n_*(*t*)) = *s*_2_, then *x_k_*(*t* + 1) = *s*_2_ with probability 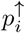. The degradation probability 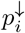 is defined similarly. SDDS can be represented as a Markov chain by specifying its transition matrix in the following way. For each variable *x_i_, i* = 1,…, *n*, the probability of changing its value is given by

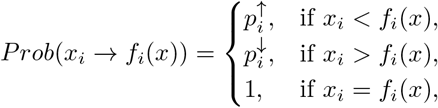

and the probability of maintaining its current value is given by

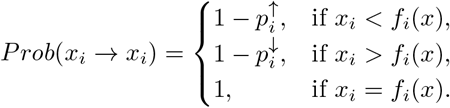

Let *x, y* ∈ *S*. The transition from *x* to *y* is given by

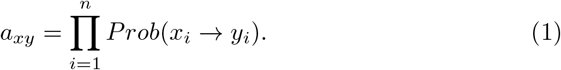

Notice that *Prob*(*x_i_* → *y_i_*) = 0 for all *y_i_* ∉ {*x_i_*, *f_i_*(*x*)}.

The stochastic framework is implemented in the toolbox to give the user with simulation options including the deterministic case. By setting all propensities equal to 1, one obtains a deterministic model. Alternatively, setting all propensities equal to 0.9 gives a 90% chance of using the regulatory function for each node and a 10% chance of keeping the current state. Likewise, setting all propensities equal to 0.5 gives a 50% chance of using the regulatory function for each node and a 50% chance of keeping the current state value. Furthermore, one could use the parameter estimation techniques for computing the propensity parameters of SDDS that have been presented in [15].

## 3 Applications: Gene expression data from experiments in axolotls

In this section we apply our method to a time-course, gene expression data that were collected during salamander (axolotls – *Ambystoma mexicanum*) tail regeneration under control and intervention conditions. Chemicals that inhibit cell-signaling activities are used as intervention agents to block tail regeneration and alter gene expression. Modeling gene interactions can provide confirmatory and novel information for developing hypotheses about the actions of cell-signaling molecules and transcription factors that orchestrate tissue regeneration.

Using our method, we generated a Boolean model for a set of 10 genes that were expressed differently during axolotl tail regeneration under control and Wnt C59 treatment, a chemical that blocks the secretion of Wnt signaling molecules from cells [16]. We note that the Wnt C59 intervention in that study inhibited tail regeneration. Seven of the genes are ligands (Areg, *Fgf9, Bmp2, Inhbb*, and *Wnt5a*) or negative feedback regulators (*Dusp6, Nradd*) of cell signaling pathways, *Sp7* is a bone-specific transcription factor, *Hapln3* is a cell adhesion molecule and *Phlda2* is an intracellular protein. We label these genes using the following variables:

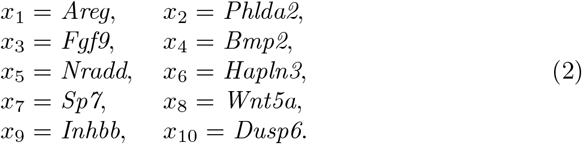

In Figure 5 we show the wiring diagrams obtained using our method for both conditions, control and intervention. These wiring diagrams present gene-by-gene interactions for the given gene expression data set. For the control case, *Wnt5a* represents a key node in the network; this Wnt signaling ligand is predicted to activate ligands that function in BMP *(Bmp2*), FGF *(Fgf9*), and TGFβ *(Inhbb*) pathways, and transcriptional regulation of bone formation (Sp7) [17], consistent with Wnt signaling playing a central, integrative role in regeneration [18] (Figure 5 a). Additionally, for the control case, we note that the well-established inhibitory effect of Dusp6 on FGF signaling is captured by this wiring diagram [19], and it also implicates *Wnt5a* as an inhibitor of *Areg* during regeneration. By comparing the intervention with the control case, we see that Wnt C59 eliminated all of the *Wnt5a* activating edges and the inhibitory edge to *Areg.* A new activating edge from *Areg* to *Nradd* suggests a novel hypothesis that Wnt C59 blockade of Wnt ligand secretion indirectly (via Nradd) inhibits the transcription of BMP and FGF pathway ligands that are required for tissue regeneration [20].

**Figure 4:**
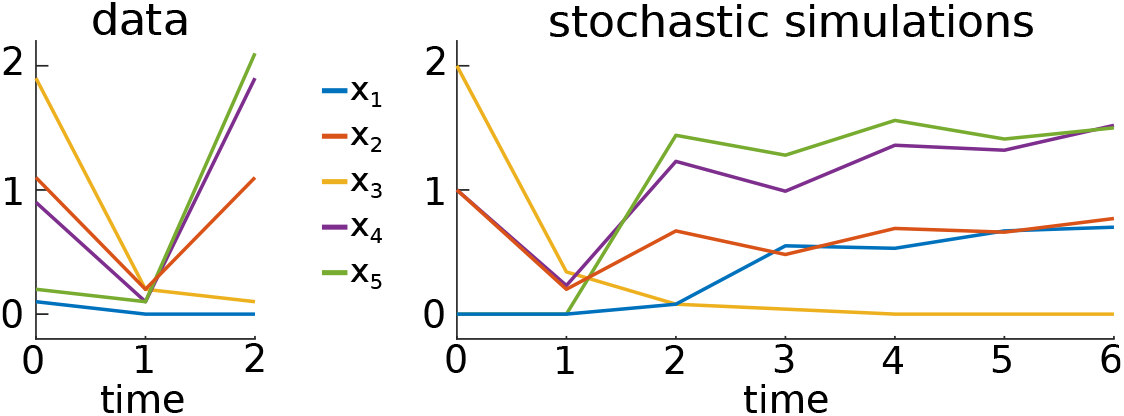
Comparison between data (only first time course shown) and stochastic simulations. Using the discretization of the initial condition of the data, 01210, we can use the model obtained to simulate the system for any arbitrary number of steps. Stochastic simulations are the average of 100 realizations, so they may take non discrete values between 0 and 2.

**Figure 5:**
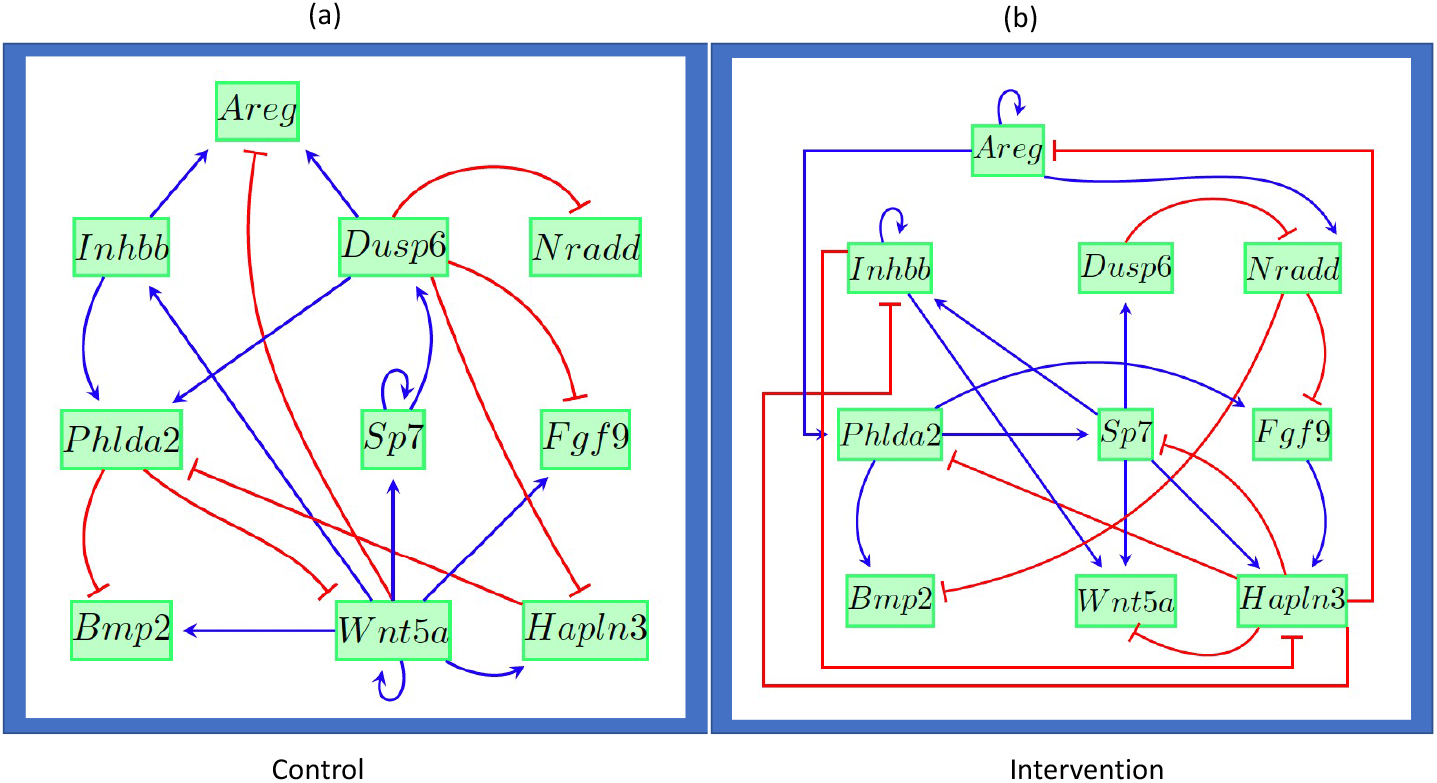
Wiring diagrams for the genes in Equation 2 for both conditions: (a) control and (b) intervention. Blue edges represent activation while red edges inhibition.

To further validate this model we compare the experimental data versus the simulations that are shown in Figures 6–7. These figures were obtained from 100 runs. Simulations using the framework SDDS [14] were performed initializing the system at the initial state 1100000001. This initialization represents a discretized version of the actual data at time 0. For the simulations in Figures 6–7 we used propensities equal to 0.9 for all variables. To assess the quality of the predictions, we use the Mean Squared Error (MSE) between the discretized data and simulated trajectories (see the Appendix for details of the MSE). We also generated simulation plots using propensities equal to 0.5 for all variables, see Figures 11–12. From the MSE values, it can be seen that the simulated trajectories with propensities equal to 0.9 give better fits of the discretized data. The propensity values can further be optimized using the method in [15].

**Figure 6:**
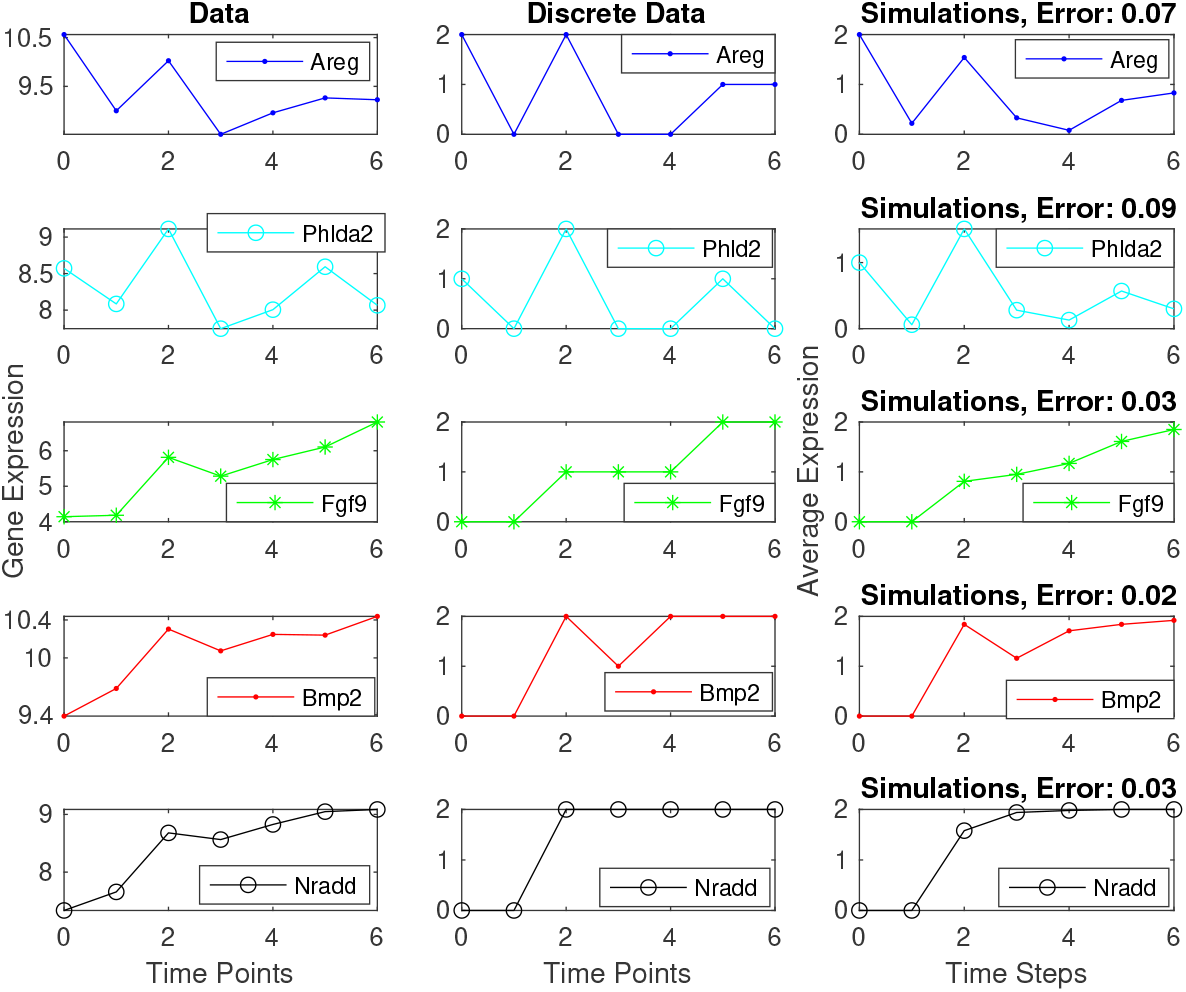
Gene expression data, discretized data, and simulations of the first five genes in Equation 2. The plots in the left panel are the experimental data, the ones in the middle are the discretized data, and the ones in the right panel are average expressions from simulations of 100 runs, all initialized at 1100000001. For the simulations, all the propensities are equal to 0.9 and the mean squared error is between the discretized data and simulated trajectories.

**Figure 7:**
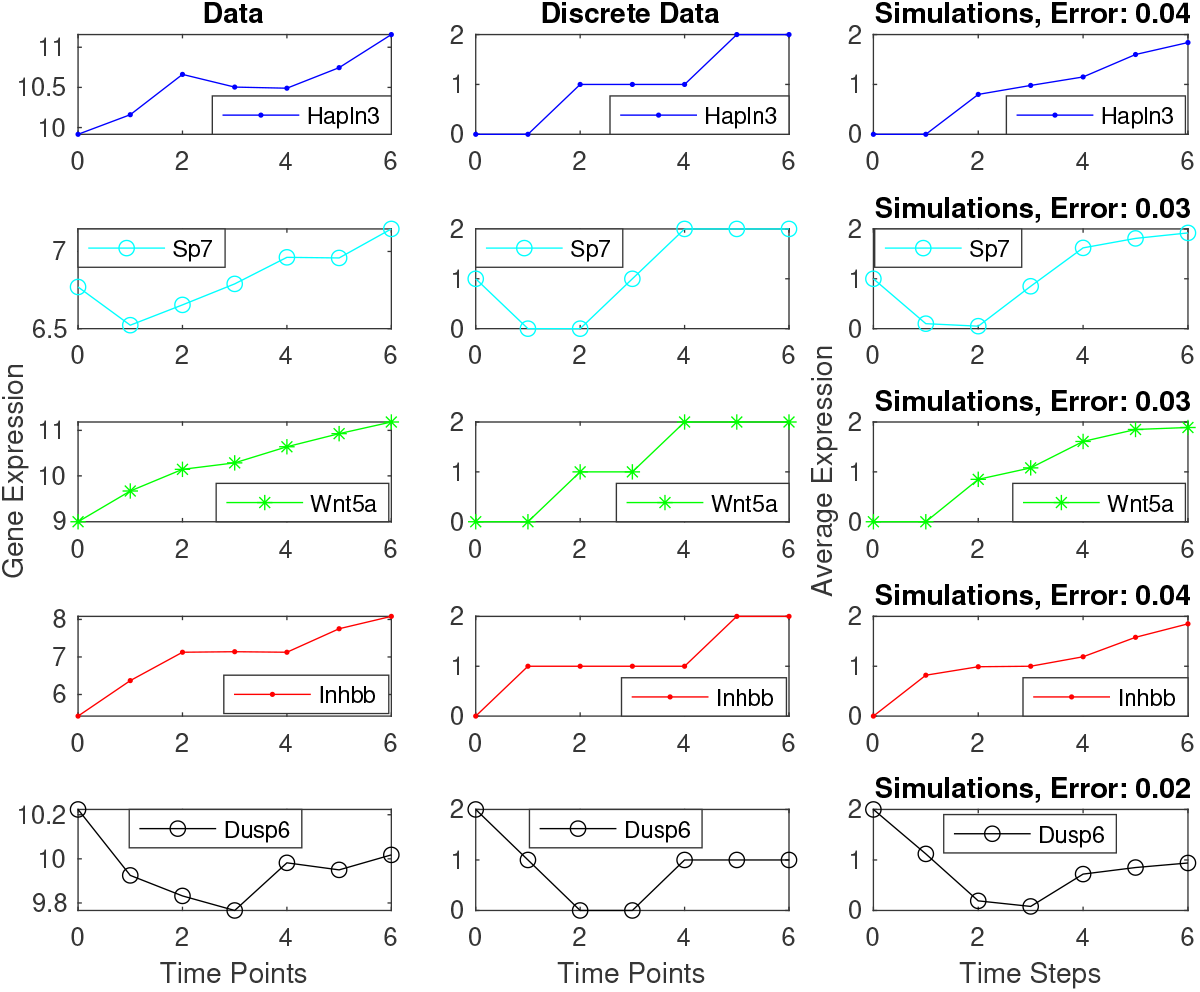
Gene expression data, discretized data, and stochastic simulations of the last five genes in Equation 2. The plots in the left panel are experimental data, the ones in the middle are discretized data, and the ones in the right panel are average expressions from simulations of 100 runs, all initialized at 1100000001. For the simulations, all the propensities are equal to 0.9 and the mean squared error is between the discretized data and simulated trajectories.

This model can further be used to attractor analysis, control, modularity, etc. However, the main result from this application is the potential novel interactions that can be experimentally tested.

## 4 Effect of data size, noise, number of levels/states, and threshold value

Since our toolbox consists in the combination of several methods/algorithms, any advantages or disadvantages of these will affect the performance of the model created. We explore some of these effects using synthetic networks so that we can compare the model constructed with the original network.

### 4.1 Effect of data size

Here we illustrate the effect of data size using a synthetic example given by the Boolean network *f* = (*f*_1_,…, *f*_5_), where

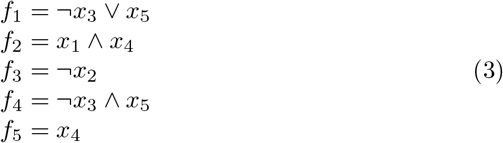

We will generate synthetic data using this Boolean network. Then, *using the data only*, we will see if the following novel trajectories can be predicted: 01111 → 10001 → 10110 → 01101 → 10000 and 00100 → 00100 (that is, 00100 is a steady state). That is, we will initialize the system at 00100 and 01111 in the model predicted by our toolbox. The results are summarized in Table 6. To illustrate the effect of the number of data points clearer, we do not consider stochasticity in this subsection (all propensities are set equal to 1).

**Table 6:** Effect of increasing the data size. *T*_1_ = {11011 → 11011}, *T*_2_ = {11010 → 11001 → 10010}, *T*_3_ = {11110 → 01001 → 10010 → 11101}, *T*_4_ = {01011 → 10011 → 11111 → 11001}. Transitions predicted correctly are indicated by bold arrows.

| Data used | Novel trajectories predicted |
| --- | --- |
| $T_1, T_2$ | $01111 \rightarrow 11011 \rightarrow 11011 \rightarrow 11011 \rightarrow 11011, 00100 \rightarrow 10000$ |
| $T_1, T_2, T_3$ | $01111 \rightarrow 01011 \rightarrow 11011 \rightarrow 11011 \rightarrow 11011, 00100 \rightarrow 00100$ |
| $T_1, T_2, T_3, T_4$ | $01111 \rightarrow 10001 \rightarrow 10110 \rightarrow 01101 \rightarrow 10000, 00100 \rightarrow 00100$ |

First, we start with the synthetic time series *T*_1_ = {11011 → 11011} and *T*_2_ = {11010 → 11001 → 10010}. With this data, our toolbox predicts that the trajectories initialized at 01111 and 00100 are 01111 → 11011 → 11011 → 11011 → 11011 and 00100 → 10000. In this case the data was not enough to recover the trajectories.

Second, we start with the synthetic time series *T*_1_, *T*_2_, and also *T*_3_ = {11110 → 01001 → 10010 → 11101}. With this data, our toolbox predicts that the trajectories initialized at 01111 and 00100 are 01111 → 01011 → 11011 → 11011 → 11011 and 00100 → 00100. In this case we see that with more data still the trajectory for 01111 was not recovered, but the model created by the toolbox correctly predicted that 11011 is a steady state.

Third, we start with the synthetic time series *T*_1_, *T*_2_, *T*_3_, and also *T*_4_ = {01011 → 10011 → 11111 → 11001}. With this data, our toolbox predicts that the trajectories initialized at 01111 and 00100 are 01111 → 10001 → 10110 → 01101 → 10000 and 00100 → 00100. That is, with the data given, the model predicted by the toolbox was able to correctly reproduce the novel trajectories. Note that *T*_1_,*T*_2_,*T*_3_,*T*_4_ represent 9 out of the 2^5^ = 32 possible transitions.

### 4.2 Effect of noise

Here we study the effect of noise in the model predicted by the toolbox. We use the Boolean network in Eq. 3 as the truth and the trajectories *T*_1_,*T*_2_,*T*_3_,*T*_4_ as data. To this data we will add noise following a uniform distribution centered at 0 and with standard deviation σ. With the noisy data, we use the toolbox to create a model and make a prediction of the trajectory with initial condition 01111.

The results are shown in Fig. 8. We see that for small noise the model is still able to make an accurate prediction of the trajectory. Since Boolean models focus on the qualitative features of he dynamics, it is robust to small noise levels. However, for large enough noise, the predicted trajectory does not match the true trajectory. A possible cause is that the first step of the discretization needs to distinguish between “low” and “high”. If noise is large, it is possible that a low value plus noise is larger than a high value plus noise and may be incorrectly discretized.

**Figure 8:**
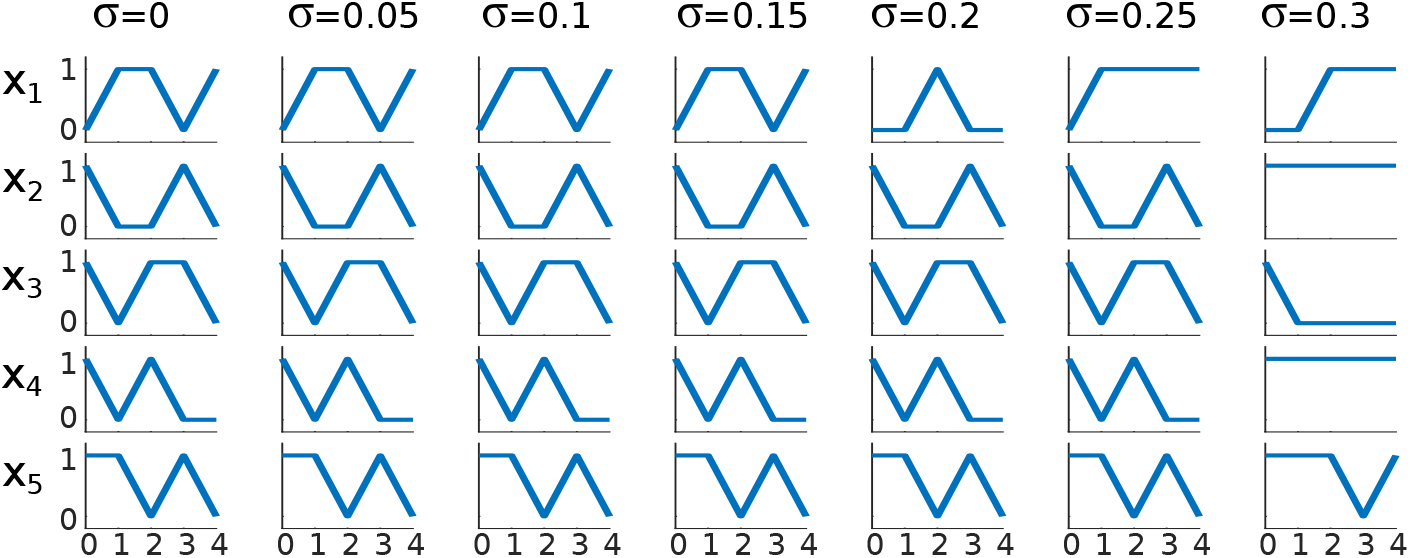
Effect of noise on predicted model. For small noise, the qualitative behavior is maintained, but as noise increases, the predicted model loses its predictive power.

### 4.3 Effect of number of levels

Here we use the network with 4 levels (or states) *f* = (*f*_1_,…, *f*_5_): {0,1, 2, 3}^5^ → {0,1, 2, 3}^4^, where

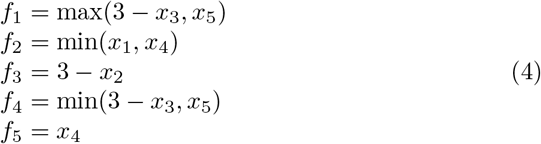

We generated 5 trajectories using this network: 32032 → 33123 → 32022 → 32122 → 22122, 23131 → 22013 → 31131 → 23213 → 31011, 22322 → 22102 → 20120 → 22302 → 20100, 03123 → 30022 → 32322 → 22102 → 20120, 03121 → 20012 → 31321 → 12202 → 20110. Using this data only, we use our toolbox to create a model and make a prediction for the novel trajectory with initial condition 03232. The true trajectory is shown in Fig. 9 (first column).

**Figure 9:**
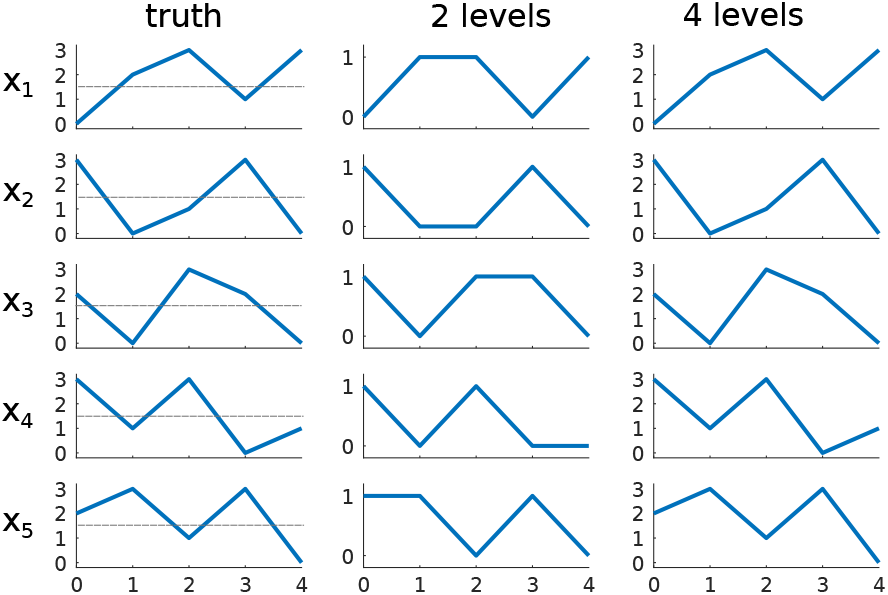
Effect of different levels. Using a coarser discretization, some features are lost. Using an appropriate number of levels, more features can be captured by the model.

Selecting a discretization of the data with 2 levels in the toolbox (0 and 1 become 0, and 2 and 3 become 1) will create a Boolean model. In this case, the initial condition 03232 would become 01111 and the Boolean trajectory will of course not match the true trajectory exactly. However, the Boolean trajectory does have the same qualitative features of the true trajectory, Fig. 9. For instance, *x*_1_ has the pattern 0 → 2 → 3 → 1 → 3 in the true trajectory. If discretized, this would be0 → 1 → 1 → 0 → 1, just like the predicted Boolean trajectory.

We then considered a discretization that included the previous one. To achieve this we considered 4 levels. Based on the data, using 4 levels is a more natural discretization and indeed, the trajectory predicted by the toolbox matches the true trajectory, Fig. 9.

### 4.4 Effect of threshold values

We now explore the effect of changing the threshold chosen for discretization on the predicted model. We use the network *f*: {0,1, 2}^3^ → {0,1, 2}^3^ given by

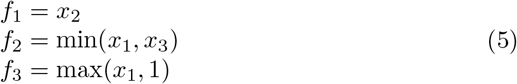

We use the data 111 → 111, 020 → 201, 002 → 001, 220 → 202 → 022 and will attempt to predict the true trajectory 020 → 201 → 012 → 101 → 011 using the model given by the toolbox. To make the comparison simpler, we choose 2 states only. Choosing different thresholds can potentially result in different models with different dynamical properties. The difference is shown in Fig. 10. Different states can be mapped to the same value if they are both less than or greater than the threshold. This causes the loss of certain features, but some coarse qualitative features are preserved.

**Figure 10:**
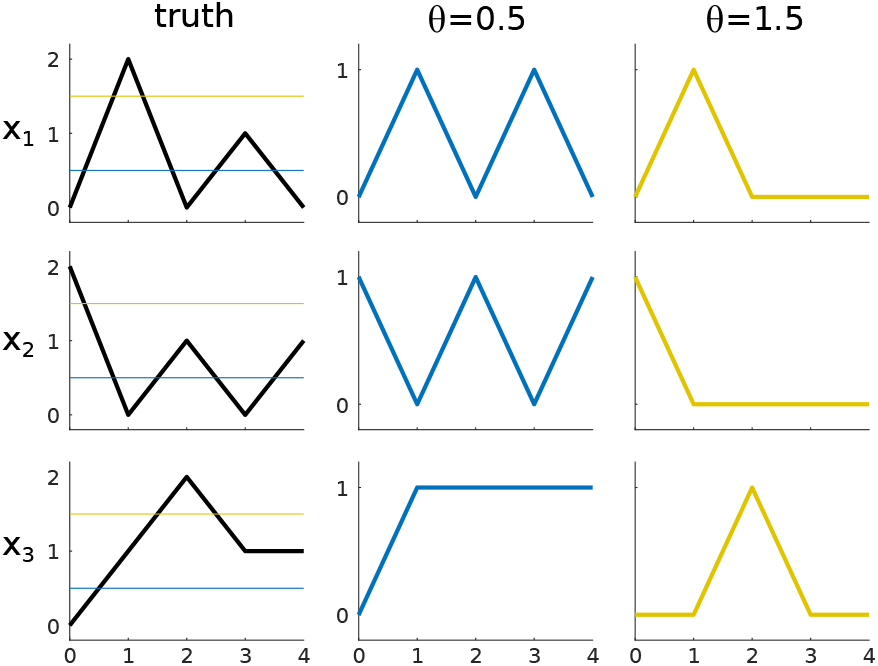
Effect of different thresholds. For thresholds around *θ* = 0.5, values that should have been 1 or 2 in the true trajectory are all mapped to 1. For thresholds around *θ* = 1.5, the values that should have been 0 or 1 are mapped to 0.

## 5 Discussion

Discrete models have been successfully used to model biological systems [8, 21]. Although several discrete modeling packages exist for their analysis (e.g., PlantSimLab [22], BoolNet [23], BNReduction [24], GinSim [25], CaSQ [26], WebMaBoSS [27]), they require an existing model or the wiring diagram to be created by the user. Few tools exist that provide an automated and easily customizable pipeline to quickly create model prototypes. Our toolbox allows the creation of model prototypes easily, which can then be used by existing modeling packages for validation, modification, or extension.

Equation learning methods in general require large amounts of data which might not be feasible in practice [12,13]. Furthermore, those approaches require knowledge of the form of the functions (some times called a library of functions) *a priori*, which may be unfeasible for unknown interactions. Even if the form of the functions is known for continuous modeling, the model obtained can be the result of parameter estimation being stuck in a local minimum. In contrast, our method can be used even with a limited number of time points. Although this does not guarantee predictive power, our toolbox does find all minimal wiring diagrams. This is important, because it can be seen as the discrete version of finding all local minima in parameter estimation for continuous models. Furthermore, our approach does not need to know the form of the functions *a priori*. We note that the discrete model resulting from our approach can be converted into a continuous model using existing approaches such as [28,29].

The limitations of our toolbox are those related to each component in the pipeline. Notably, if the discretization considers two instances of the same value as different due to noise (e.g. 0.7 as 0 and 0.9 as 1), this can cause overfitting. Selecting the correct number of levels of the model is also important and can cause missing some features if the number of levels is too low or overfitting if the number of levels is too large. Also, the selection of thresholds can make a difference on which features of the true dynamics are correctly predicted with the inferred model. Another limitation is that it is not known how much data is needed to guarantee that the predicted model is “close” to the true network unless the network is known *a priori.*

For the purpose of reproducibility, we provide all the data and the code that we use in our toy example and application which can be accessed through this link: github.com/alanavc/prototype-model.

## A Code Usage

Here we show how the Matlab code is used. All code files and examples can be found at Github github.com/alanavc/prototype-model. The toy example and application are included.

The input files must be in the form of trajectories (one line per state of the system). Different trajectories must be in different files and all trajectories files must have the same prefix in the name. For example, `timecourse1.txt, timecourse2.txt, timecourse3.txt, timecourse4.txt`, etc. For our example in Methods, the content of timecourse1.txt would be the following.

~~~
0.1,1.1,1.9,0.9,0.2
0.0,0.2,0.2,0.1,0.1
0.0,1.1,0.1,1.9,2.1
~~~

Once we are in the folder `codes`, we can discretize the data. Prior to this we need to specify the prefix for the time course files and the file where the data will be saved.

~~~
> clear all
> time_course_prefix=‘timecourse’;
> num_levels=[3,2,3,3,3];
> Íile_for_disc_data=‘data0.mat’;
> % discretize data
> addpath(‘discretize_data’)
> discretize(time_course_prefix,num_levels,file_for_disc_data)
~~~

Then, we generate all minimal wiring diagrams using the Matlab function `generate_wiring_diagrams`. Prior to this, we need to specify the file where we will save all wiring diagrams.

~~~
> %create wiring diagrams
> file_for_wiring_diagrams=‘WD0.mat’;
> addpath(‘create_WD’)
> generate_wiring_diagrams(file_for_disc_data,file_for_wiring_diagrams)
~~~

To select the best wiring diagram we use the command `select_best_wiring_diagram` as follows.

~~~
> %select wiring diagram
> addpath(‘select_WD’)
> W=select_best_wiring_diagram(file_for_wiring_diagrams);
~~~

The variable `W` is a matrix that contains the best wiring diagram that fits the data. The user can modify this to account for prior knowledge or to try any other of the wiring diagrams (found in file `WD0.mat`) for exploration.

We use the function `generate_monotone_functions` to create the model that fits the data and has the wiring diagram given. The model will be saved as a truth table for each variable.

~~~
> %create model
> í?le_íor_model=‘model0.mat’;
> addpath(‘create_Mode?)
> generate_monotone_functions(time_course_prefix,W,file_for_model)
~~~

To simulate the model we use the following code.

~~~
> %simulate model
> addpath(‘simulate_mode?)
> init_state = [01210];
> num_simulations = 100;
> num_steps = 6;
> num_vars = 5;
> % Propensity matrix
> propensity_matrix = 0.8*ones(2,num_vars);
> mean_trajectories = simulate(file_for_model,propensity_matrix, init_state,num_steps,num_simulations);
~~~

The mean trajectories are saved in `mean_trajectories` and can be plotted using standard Matlab commands.

## B Inferred Model

Here we describe the inferred models under the control and intervention conditions that we described in the Applications Section. We used *n* =10 genes for both cases. The number of input variables in each discrete function in the *control* case is given in the following vector:

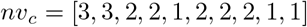

And the number of inputs in each discrete function for the *intervention* case is given in the following vector:

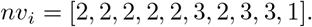

The threshold for selecting the wiring diagram for the control case is *τ_c_* = 1/2 while for the intervention case is *τ_i_* = 3/4. The maximum number of inputs for both cases (control and intervention) is *m* = 3. In Table 7 we list the input variables for each gene for the control case. This table is a m-by-n matrix. Since the number of variables may vary between different functions, only the first *nv_c_*(*i*) elements are relevant in the *i^th^* column of Table 7. The remaining spots in each column are set to −1 so that the whole table can be encoded as a matrix. Similarly for the intervention case, we list the input variables for each gene in Table 8.

The number of levels for the discretization in both cases is *p* = 3. In Table 9 we provide the truth table for the control case in compact form. Similarly, the truth table for the intervention case is given in Table 10. These tables have size *p^m^*-by-*n*. Since the length of the truth tables may vary between different functions, only the first *p*^*nv*(*i*)^ bits are relevant in the *i^th^* column of Table 9. The remaining spots in each column are set to −1 so that the whole table can be encoded as a matrix. The entries of the columns of *F* are ordered so that they correspond to the binary input arrays in lexicographic order. For example, the first column in Table 10 with relevant entries 1,0,0,1,1,0,2,2,1 represents the truth table that satisfies *h*(0,0) = 1, *h*(0,1) = 0, *h*(0, 2) = 0, *h*(1,0) = 1, *h*(1,1) = 1, *h*(1, 2) = 0, *h*(2,0) = 2, *h*(2,1) = 2, and *h*(2, 2) = 1.

**Table 7:** Input variables for the control case. Each column gives the inputs for each gene in Eq. 2.

| $x_1$ | $x_2$ | $x_3$ | $x_4$ | $x_5$ | $x_6$ | $x_7$ | $x_8$ | $x_9$ | $x_{10}$ |
| --- | --- | --- | --- | --- | --- | --- | --- | --- | --- |
| 8 | 6 | 8 | 2 | 10 | 8 | 7 | 2 | 8 | 7 |
| 9 | 9 | 10 | 8 | -1 | 10 | 8 | 8 | -1 | -1 |
| 10 | 10 | -1 | -1 | -1 | -1 | -1 | -1 | -1 | -1 |

**Table 8:** Input variables for the intervention case. Each column gives the inputs for each gene in Eq. 2.

| $x_1$ | $x_2$ | $x_3$ | $x_4$ | $x_5$ | $x_6$ | $x_7$ | $x_8$ | $x_9$ | $x_{10}$ |
| --- | --- | --- | --- | --- | --- | --- | --- | --- | --- |
| 1 | 1 | 2 | 2 | 1 | 3 | 2 | 6 | 6 | 7 |
| 6 | 6 | 5 | 5 | 10 | 7 | 6 | 7 | 7 | -1 |
| -1 | -1 | -1 | -1 | -1 | 9 | -1 | 9 | 9 | -1 |

Finally, the propensities that we used for the simulations in Figures 11–12 are all equal to 0.9. That is, 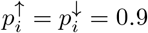 for all *i* = 1,…, 10.

## C Simulations with Different Propensities

In the main text we used propensities equal to 0.9 for the simulations. Here we show simulations results when all the propensities are equal to 0.5. That is, 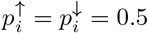 for all *i* = 1,…, 10.

## D Mean Squared Error

Let *Y* is the vector of discretized data and *Ŷ* is a vector of average expressions from simulated trajectories. If *Y* and *Ŷ* have the same length, we calculate the Mean Squared Error (MSE) of the predictor (or the Mean Squared Prediction Error) using the following formula:

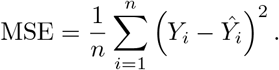

Note that the MSE is the “mean” 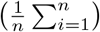 of the “squares of the errors” 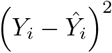.

**Table 9:** Truth tables (in compact form) for the control case in the Applications Section.

| $x_1$ | $x_2$ | $x_3$ | $x_4$ | $x_5$ | $x_6$ | $x_7$ | $x_8$ | $x_9$ | $x_{10}$ |
| --- | --- | --- | --- | --- | --- | --- | --- | --- | --- |
| 0 | 0 | 1 | 2 | 2 | 1 | 0 | 1 | 1 | 0 |
| 0 | 0 | 1 | 2 | 2 | 1 | 1 | 2 | 1 | 1 |
| 0 | 0 | 0 | 2 | 0 | 0 | 1 | 2 | 2 | 1 |
| 0 | 0 | 1 | 0 | -1 | 1 | 0 | 0 | -1 | -1 |
| 2 | 2 | 1 | 1 | -1 | 1 | 2 | 1 | -1 | -1 |
| 2 | 2 | 0 | 2 | -1 | 0 | 2 | 2 | -1 | -1 |
| 0 | 0 | 2 | 0 | -1 | 2 | 0 | 0 | -1 | -1 |
| 2 | 2 | 2 | 1 | -1 | 2 | 2 | 1 | -1 | -1 |
| 2 | 2 | 0 | 1 | -1 | 0 | 2 | 1 | -1 | -1 |
| 0 | 0 | -1 | -1 | -1 | -1 | -1 | -1 | -1 | -1 |
| 0 | 0 | -1 | -1 | -1 | -1 | -1 | -1 | -1 | -1 |
| 0 | 0 | -1 | -1 | -1 | -1 | -1 | -1 | -1 | -1 |
| 0 | 0 | -1 | -1 | -1 | -1 | -1 | -1 | -1 | -1 |
| 1 | 1 | -1 | -1 | -1 | -1 | -1 | -1 | -1 | -1 |
| 1 | 1 | -1 | -1 | -1 | -1 | -1 | -1 | -1 | -1 |
| 0 | 0 | -1 | -1 | -1 | -1 | -1 | -1 | -1 | -1 |
| 1 | 1 | -1 | -1 | -1 | -1 | -1 | -1 | -1 | -1 |
| 1 | 1 | -1 | -1 | -1 | -1 | -1 | -1 | -1 | -1 |
| 0 | 0 | -1 | -1 | -1 | -1 | -1 | -1 | -1 | -1 |
| 0 | 0 | -1 | -1 | -1 | -1 | -1 | -1 | -1 | -1 |
| 0 | 0 | -1 | -1 | -1 | -1 | -1 | -1 | -1 | -1 |
| 0 | 0 | -1 | -1 | -1 | -1 | -1 | -1 | -1 | -1 |
| 1 | 0 | -1 | -1 | -1 | -1 | -1 | -1 | -1 | -1 |
| 1 | 0 | -1 | -1 | -1 | -1 | -1 | -1 | -1 | -1 |
| 0 | 0 | -1 | -1 | -1 | -1 | -1 | -1 | -1 | -1 |
| 1 | 0 | -1 | -1 | -1 | -1 | -1 | -1 | -1 | -1 |
| 1 | 0 | -1 | -1 | -1 | -1 | -1 | -1 | -1 | -1 |

**Table 10:** Truth tables (in compact form) for the intervention case in the Applications Section.

| $x_1$ | $x_2$ | $x_3$ | $x_4$ | $x_5$ | $x_6$ | $x_7$ | $x_8$ | $x_9$ | $x_{10}$ |
| --- | --- | --- | --- | --- | --- | --- | --- | --- | --- |
| 1 | 1 | 1 | 0 | 0 | 0 | 0 | 1 | 1 | 0 |
| 0 | 0 | 0 | 0 | 0 | 0 | 0 | 2 | 2 | 0 |
| 0 | 0 | 0 | 0 | 0 | 0 | 0 | 2 | 2 | 1 |
| 1 | 1 | 2 | 1 | 2 | 0 | 1 | 1 | 1 | -1 |
| 1 | 1 | 0 | 0 | 0 | 0 | 1 | 2 | 2 | -1 |
| 0 | 0 | 0 | 0 | 0 | 0 | 0 | 2 | 2 | -1 |
| 2 | 2 | 2 | 2 | 2 | 2 | 2 | 1 | 1 | -1 |
| 2 | 2 | 2 | 2 | 2 | 0 | 2 | 2 | 2 | -1 |
| 1 | 2 | 0 | 0 | 1 | 0 | 0 | 2 | 2 | -1 |
| -1 | -1 | -1 | -1 | -1 | 1 | -1 | 0 | 0 | -1 |
| -1 | -1 | -1 | -1 | -1 | 0 | -1 | 2 | 2 | -1 |
| -1 | -1 | -1 | -1 | -1 | 0 | -1 | 2 | 2 | -1 |
| -1 | -1 | -1 | -1 | -1 | 1 | -1 | 0 | 0 | -1 |
| -1 | -1 | -1 | -1 | -1 | 0 | -1 | 2 | 2 | -1 |
| -1 | -1 | -1 | -1 | -1 | 0 | -1 | 2 | 2 | -1 |
| -1 | -1 | -1 | -1 | -1 | 2 | -1 | 1 | 1 | -1 |
| -1 | -1 | -1 | -1 | -1 | 0 | -1 | 2 | 2 | -1 |
| -1 | -1 | -1 | -1 | -1 | 0 | -1 | 2 | 2 | -1 |
| -1 | -1 | -1 | -1 | -1 | 1 | -1 | 0 | 0 | -1 |
| -1 | -1 | -1 | -1 | -1 | 1 | -1 | 0 | 0 | -1 |
| -1 | -1 | -1 | -1 | -1 | 0 | -1 | 0 | 0 | -1 |
| -1 | -1 | -1 | -1 | -1 | 1 | -1 | 0 | 0 | -1 |
| -1 | -1 | -1 | -1 | -1 | 1 | -1 | 0 | 0 | -1 |
| -1 | -1 | -1 | -1 | -1 | 0 | -1 | 0 | 0 | -1 |
| -1 | -1 | -1 | -1 | -1 | 2 | -1 | 0 | 0 | -1 |
| -1 | -1 | -1 | -1 | -1 | 2 | -1 | 1 | 1 | -1 |
| -1 | -1 | -1 | -1 | -1 | 0 | -1 | 1 | 1 | -1 |

**Figure 11:**
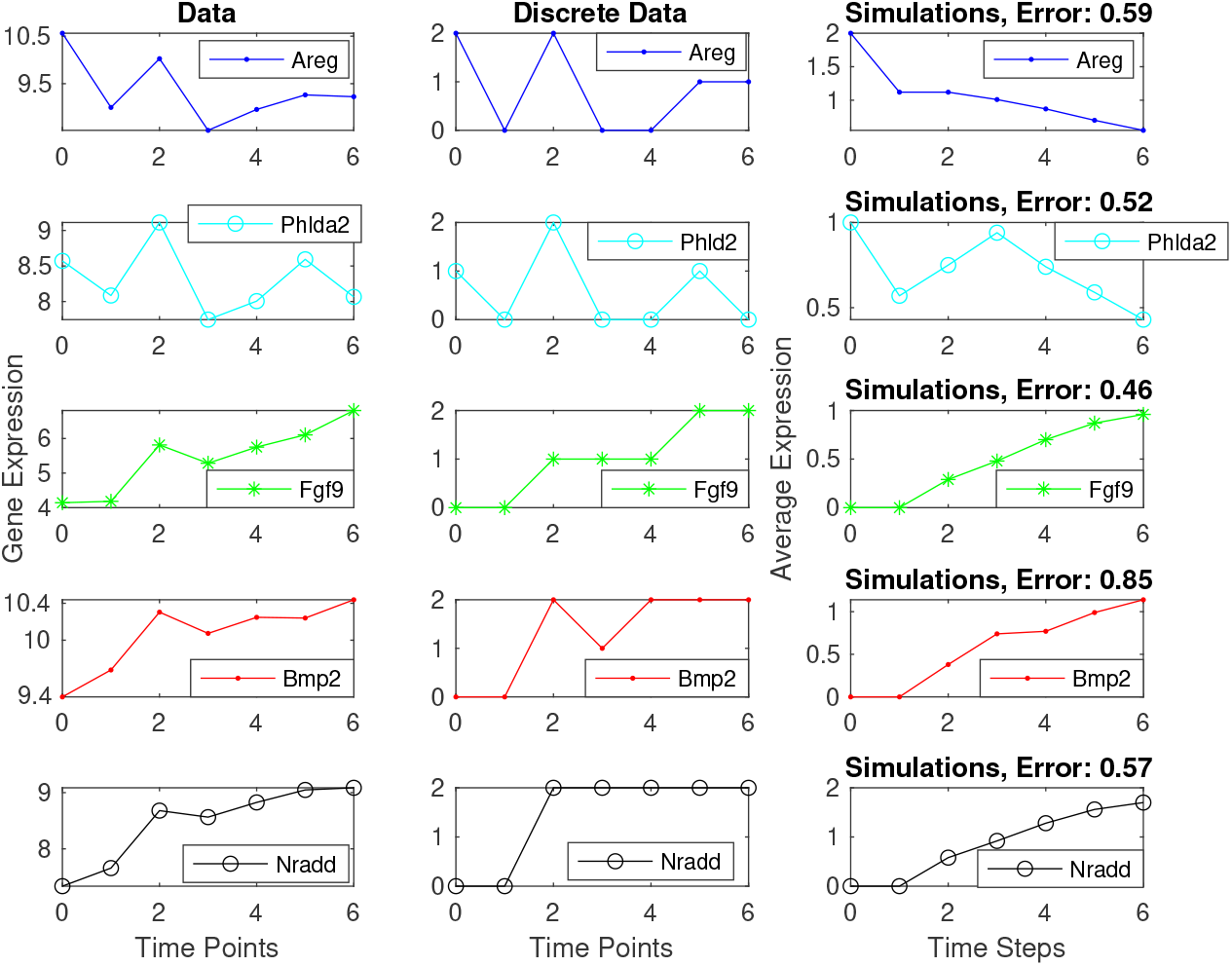
Gene expression data, discretized data, and simulations of the first five genes in Equation 2. The plots in the left panel are the experimental data, the ones in the middle are the discretized data, and the ones in the right panel are average expressions from simulations of 100 runs, all initialized at 1100000001. For the simulations, all the propensities are equal to 0.5 and the mean squared error is between the discretized data and simulated trajectories.

**Figure 12:**
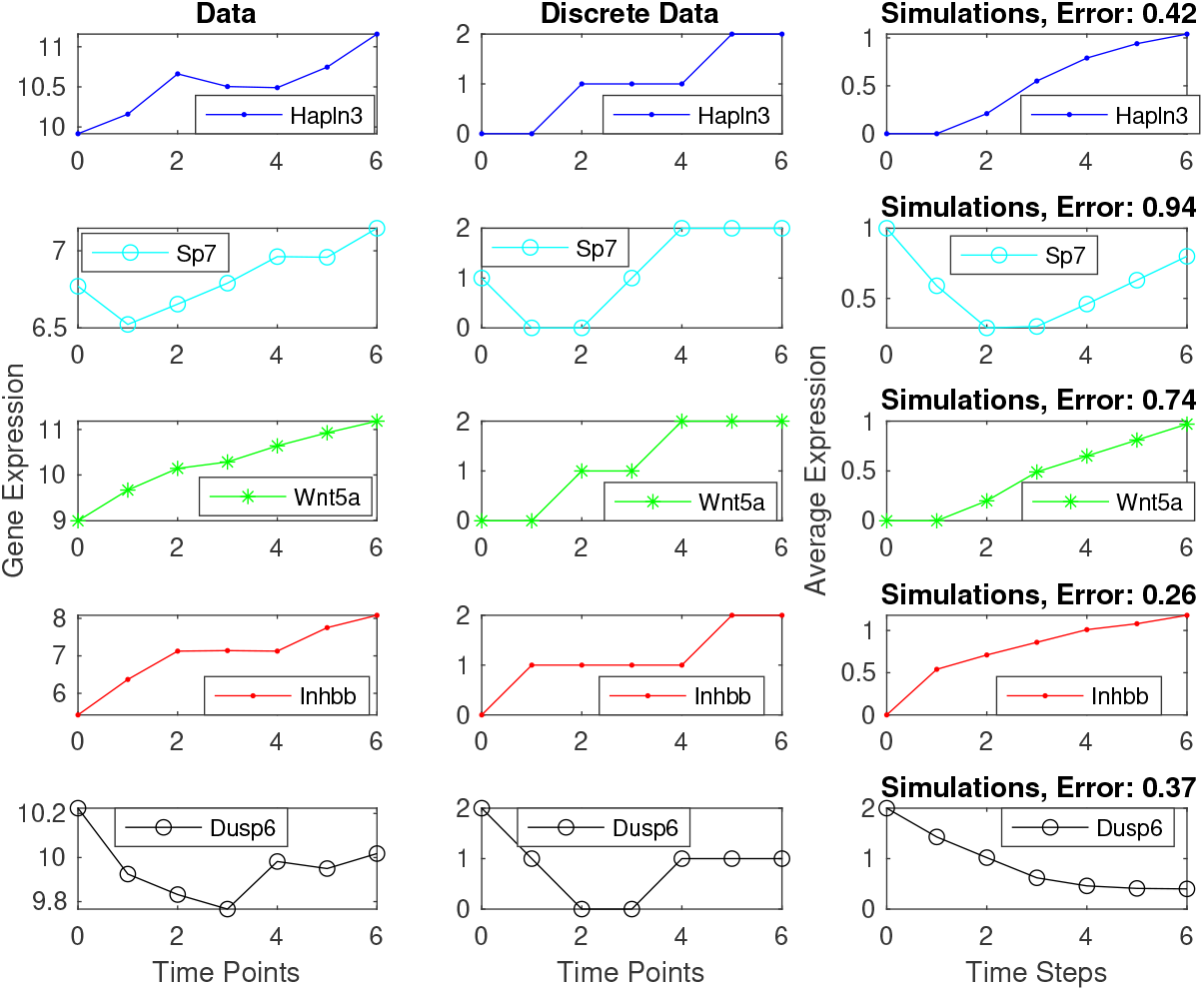
Gene expression data, discretized data, and stochastic simulations of the last five genes in Equation 2. The plots in the left panel are experimental data, the ones in the middle are discretized data, and the ones in the right panel are average expressions from simulations of 100 runs, all initialized at 1100000001. For the simulations, all the propensities are equal to 0.5 and the mean squared error is between the discretized data and simulated trajectories.

